# Using passive telemetry and environmental variables to predict Silver Carp (*Hypophthalmichthys molitrix)* movement cues on the northwestern edge of their invasion front

**DOI:** 10.1101/2023.12.07.569972

**Authors:** Lindsey A. P. LaBrie, Jeff S. Wesner

## Abstract

Silver carp *Hypophthalmichthys molitrix*, are a highly mobile aquatic invasive species in the United States. The James River, South Dakota, USA, is a tributary of the Missouri River and is considered the northwestern leading edge of their invasion front. Understanding silver carp movement patterns on the cusp of this invasion is key to combatting the northwestward expansion of the species. We used passive telemetry to observe large-scale movement patterns of silver carp in the James River, South Dakota. Fifty silver carp were implanted with acoustic transmitter tags in June 2021, and movement data was recorded over a 1.5-year period. Most individuals exhibited site fidelity and stayed within the James River throughout the duration of the study. We used environmental data (i.e., temperature, dissolved oxygen, daily mean discharge, the change in discharge over 24 h and 48 h) and movement data collected from passive telemetry receivers to understand and predict silver carp movement cues in the James River. Daily mean discharge (“flow”) was the most important predictor of silver carp movement in the James River. As flow increased, the probability of movement increased from 59% (95% CrI: 34% to 81%) at 1.5 m^3^/s to 94% (95% CrI: 80 to 99%) at 100 m^3^/s. In this study, silver carp exhibited a high propensity for movement within the James River, especially during periods of high flow. To prevent further northwestward expansion of these fish, silver carp movements must continue to be monitored and removal and prevention of further range expansion must be prioritized.

## Introduction

Aquatic invasive species are a major threat to freshwater ecosystems globally due to their increased ability to outcompete native species, alter food webs, or change the physical conditions of the invaded ecosystem (Brown and Moyle 1991; Habit et al. 2010). Human activities are a major pathway for invasive species introductions in aquatic environments (i.e. intentional releases, ballast water transfers) (Ricciardi and Rasmussen 1998), though only very few introduced species can or will survive long enough in a new environment to become invasive (Ricciardi 2013). Invasions are often exacerbated by human activities, allowing invaders to quickly become established in systems that have high anthropogenic modifications (Moyle and Light 1996). A key component to the invasion potential of a species is its ability to disperse and successfully colonize new habitats (Kolar and Lodge 2001).

Silver carp *Hypophthalmichthys molitrix,* were first brought to the United States and introduced into aquaculture ponds and wastewater treatment facilities in Arkansas in the early 1970s as a means of biocontrol of algae and to increase water clarity (Kolar et al. 2005). Soon after their introduction, they escaped containment and began establishing populations throughout the Mississippi River Basin and beyond (Nico et al. 2023) and are now present in many rivers throughout the midwestern and southern United States. Silver carp arrived in rivers of the midwestern United States in the early to mid 1990s (Kolar et al. 2005). Silver carp movement patterns have been well-documented in the United States in rivers such as the Wabash (Coulter et al., 2016; Coulter et al., 2022), Illinois (Coulter et al. 2018; DeGrandchamp 2006; DeGrandchamp et al. 2008), and Mississippi (Calkins et al. 2012; Fritts et al. 2021; Vallazza et al. 2021). A combination of biological and environmental cues influence silver carp movements in other well-studied systems. Silver carp have been observed embarking on large upstream movements when stream discharge and stream temperature increase, especially in the Spring and following large increases in stream discharge (Coulter et al. 2016; Coulter et al. 2018; DeGrandchamp et al. 2008). However, much of the existing movement research focuses on the prevention of fish passage into the Great Lakes ecosystems (Chapman et al. 2021; Cooke and Hill 2010; Cuddington et al. 2014; Parker et al. 2016) on the northeastern part of their expanding range. Movement research is currently limited in the Missouri River Basin and its tributaries.

Streams in eastern South Dakota are on the northwestern-most leading edge of the silver carp invasion front (Hayer et al. 2014), and silver carp were first documented in portions of the Missouri River in South Dakota in 2003 (Kolar et al. 2005). A major barrier to upstream fish passage, Gavin’s Point Dam on the mainstem Missouri River near Yankton, South Dakota has thus far prevented movement of silver carp into the upstream reservoirs and sections of the Missouri River. However, silver carp can freely swim between the mainstem Missouri and its tributaries downstream of Gavin’s Point Dam.

The James River, a major tributary to the Missouri River in eastern South Dakota, is a highly human-modified system, with at least 230 low-head dams and impoundments throughout (Shearer and Berry Jr 2003). The dams and impoundments provide some resistance to further upstream movement of these and other species of fish, including native migratory species such as the blue sucker *Cycleptus elongatus,* (Carlson 2022) but silver carp have surpassed these smaller barriers and have been recorded at locations as far upstream as Jamestown, North Dakota in the James River (Nico et al. 2023). The environmental factors that trigger silver carp movements and the overall movement patterns of silver carp have not previously been studied in the James River. It has also never been determined if the silver carp populations within the James River are resident populations, or if they primarily reside within the mainstem Missouri River and shift to the tributaries during different times of year in accordance with spawning runs or resource availability.

The purpose of this study is to determine which environmental factors influence silver carp movement throughout the James River and to determine whether the tagged population resides within the James River intermittently or continuously. Understanding silver carp movement cues and habitat preferences within this Missouri River tributary should ultimately help elucidate ways that this species can be effectively managed in South Dakota rivers and in similar river systems on this side of the invasion front.

## Methods

### Study Organism and Study Site

The James River begins in eastern North Dakota and flows south through the entirety of South Dakota to its confluence with the Missouri River. The James River is the largest tributary to the Missouri River between Gavin’s Point Dam near Yankton, South Dakota, and the confluence of the Platte and Missouri Rivers on the Nebraska-Missouri border (Muth and Schmulbach 1984). It is highly sinuous, traveling a downstream distance of 1,143 river kilometers (rkm), but only covering a straight distance of 400 km (Carreiro 2000). The landscape is historically categorized by glaciated drift prairie, but municipalities and farmland are the contemporary predominant features on the landscape (Bartlett 1984). There are at least 230 low-head dams on the James River (Shearer and Berry Jr 2003), which act as potential barriers to fish movement in times of low water but become submerged during high flow events and allow for fish passage (Carlson 2022).

### Acoustic tag deployment

We used passive telemetry to record silver carp movement data in the James River. Silver carp were captured using boat electrofishing on a Smith-Root GPP 7.5 electrofishing boat (pulsed DC current; 120 Hz; 11-14 A; 170 V; 40-60% power) (Carlson 2022), which also served as elecroanesthesia prior to surgery. The total length (mm) and weight (g) of each individual was recorded upon capture. We ensured that the tag weight represented <2% of the total mass of each fish prior to insertion (Winter 1983). A one-and-a-half to two-inch incision was made on the ventral side of each fish between the pelvic and pectoral fins, and an individually identifying InnovaSea V16 transponding tag (model V16-4H; 69kHz; ping rate: 30 - 60 seconds; tag life: 1328 days; 16 mm diameter, 68 mm length, 24 g weight; henceforth: V16 tag) (Bedford, Nova Scotia, Canada) was inserted into the body cavity. Sterile chromic catgut sutures were used to make three simple interrupted sutures postoperatively. Tools were sterilized with 50 percent chlorhexidine between procedures, and gloves were worn during the procedures and exchanged between surgeries. Each surgery took an average of five minutes to complete. Post-operation, the fish were monitored to ensure they were swimming upright in the live well before being released back into the James River in a location ≤ 1 river kilometer (rkm) away from the original point of capture. Capture of individuals and implantation surgeries were performed according to the University of South Dakota’s Institutional Animal Care and Use Committee protocol (#10-08- 20-23D).

### Receiver deployment

Downstream visibility distance (or line-of-sight distance) and bridge piling shape were two of the biggest considerations for receiver deployment. During the summer of 2021, we added eleven InnovaSea model Vr2Tx (n=5) and Vr2W (n=6) 69kHz (InnovaSea, Bedford, Nova Scotia) passive telemetry receivers to the existing telemetry network in the James River for a total of 21 receivers spanning from rkm 1 to rkm 358. Receivers were mounted in one of two setup designs: pipe-mounted or frame-mounted, as described in Carlson et al. (2023). The receivers in pipe mounts were deployed in a 3.05 m long, 10 cm diameter polyvinyl chloride (PVC) pipe on the downstream side of round or square bridge pilings. Each receiver was suspended from inside the top of the PVC pipe with two ∼3 m long pieces of 6 mm gage wire so that the inverted receiver was exposed out of the bottom of the pipe by ∼5 cm. (Figure S1a). In-stream frame mounts were constructed out of 2 cm diameter iron bars in either a triangular or square cage shape (Figure S1b and S1c) and secured to an onshore anchor (such as a tree) or nearby bridge piling (Figures S2a and S2b).

Data was offloaded from each receiver two to four times during the duration of the study. Batteries were replaced two times throughout the study, and desiccant packets were replaced at each battery change to prevent moisture from building within the receiver casing. Any biofouling that had accumulated on the receiver was removed to prevent detection interference. Three receivers were moved during the study period. Of the three, two were switched from bridge mounts to frame mounts due to silting in and low water levels, and one was removed entirely from the network due to a hardware malfunction. To collect habitat data, four HOBO temperature loggers, one dissolved oxygen (DO) sensor, and one depth logger were placed intermittently along the telemetry network (Figure 1a). Discharge data were downloaded from USGS gage number 06478500 (Scotland, SD), using the *dataRetrieval* package (De Cicco and Hirsch 2014) in Program R (version 4.1.1; R Core Team 2022).

**Figure 1.**
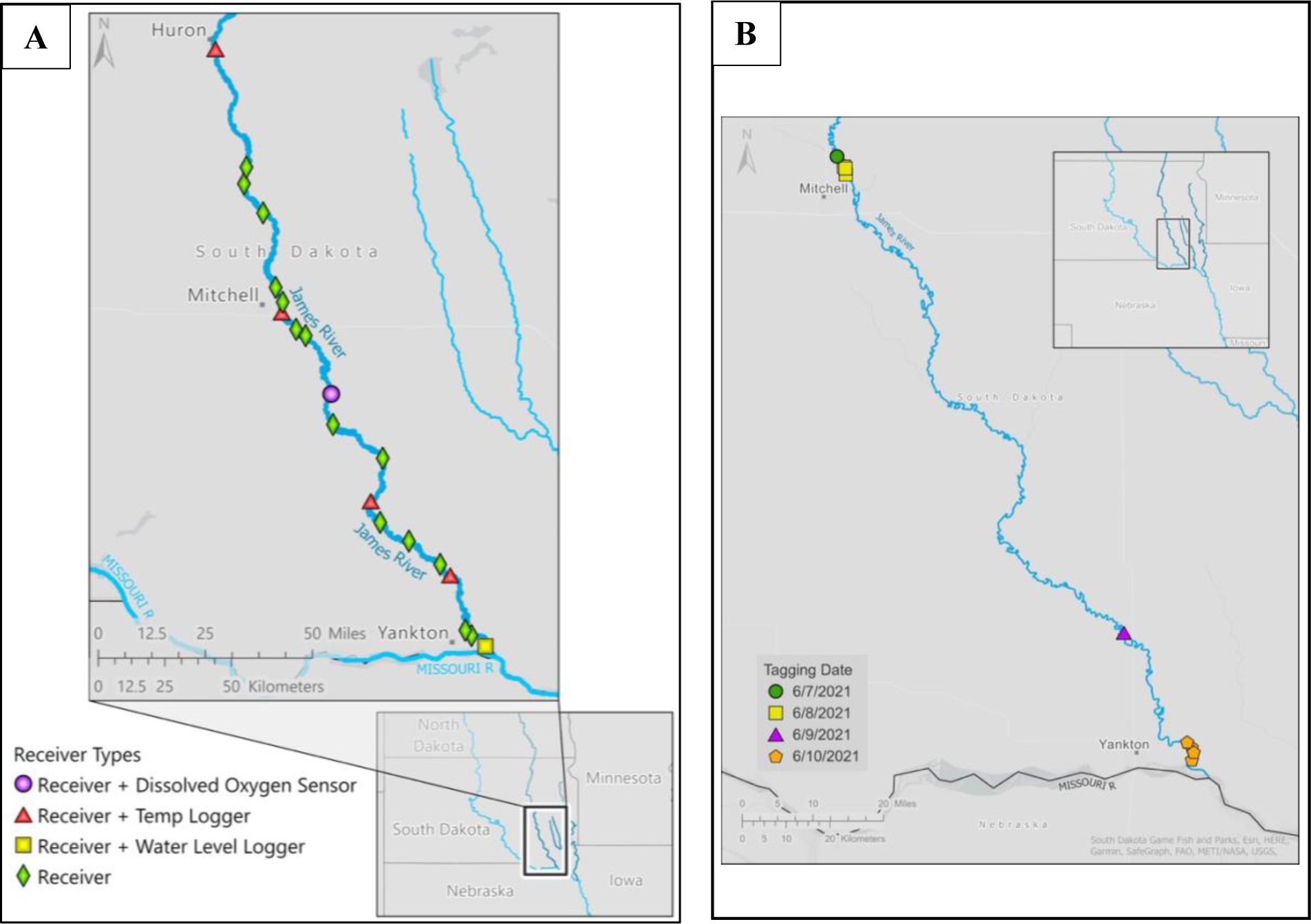
a): Map of receiver and environmental sensor locations in the James River, SD with an inset map depicting the larger area for context; 1b) silver carp release points in the James River, SD, from June 7 through June 10, 2021, with a small inset map depicting the larger area for context. Note the different map scales.

### Range test

We conducted a range test of our transmitters in mid to late summer 2022 to evaluate detection probabilities at various downstream distances from the receiver. We performed the range test using a V16 tag with the same specifications as our surgically implanted fish, except that the ping rate was set to 90 second intervals with no delay. We found that detection probability was almost 100% at 100 m downstream of the receiver. The downstream line-of-sight distance at each receiver on the James River was at least 100m (Carlson et al. 2023), and we are confident that any fish swimming near or past a receiver were detected.

### Data analysis

We kept the first instance of detection for each fish at each receiver on each day that it was detected and filtered out any subsequent data from that day unless the fish was detected on an additional receiver in the same day, in which case a datapoint was kept from the subsequent receiver following the same rule.

We used the *brms* package (Bürkner 2017) in Program R (version 4.1.1; R Core Team 2022) to create models to predict whether daily mean discharge (“flow”), a change in discharge over 24 and/or 48 hours, average weekly temperature, average weekly DO or a combination of these environmental factors best predicted silver carp movement in the James River. We fit a total of eighteen Bayesian linear models using different combinations of these variables. We estimated the probability of movement using a Bernoulli likelihood with a logit link using the following equation:

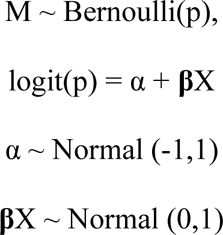

Where M indicates whether a fish moved (M=1) or not (0), p is the probability of movement versus non-movement, α is the intercept, **β**X is a linear equation representing up to five predictor variables from the six observed parameters (daily mean flow in cubic feet per second, change in discharge over 24h, change in discharge over 48h, weekly mean temp, weekly mean DO). All models were run with four chains and 2000 iterations. Then we chose the best fitting model using WAIC (Hooten & Hobbs 2015) (Table 2) and visualized the posterior predictions using *ggplot* (Wickham et al. 2016). We ran a sensitivity analysis on the best fitting model to assess the impact of the priors by doubling the standard deviation values to Normal (-1,2) for the intercept and Normal (0,2) for the beta values.

## Results

Fifty silver carp were implanted with InnovaSea V16 acoustic tags from June 7 to June 10, 2021. Fish had a mean weight of 3,019 g (range: 1042 g to 5454 g) and mean total length of 625 mm (range: 452 mm to 777 mm) (Table 1). A total of 25 transmitters were implanted near Mitchell, SD (43.760925°, -97.988167°), 11 transmitters were implanted northwest of Yankton, SD, near the Schramm public boat access (43.057850°, -97.398710°), and the final 14 transmitters were implanted approximately 2 rkm upstream of the confluence of the James and Missouri rivers (42.880131°, -97.279514°) (Figure 1b). Individual sex was not recorded. The discharge within the James River was relatively low at the time of tagging and there appeared to be little to no evidence of fallback occurrences from tagging (fish drifting downstream due to effects or complications from surgery), so we did not exclude any movement data from our analysis. However, fifteen fish were not detected in the receiver network after the date of tagging and thus had an “undetermined” fate and were not included in any analysis. Four additional fish were detected only on one receiver during multiple different dates (net recorded movement = 0) but were included in the analyses.

**Table 1:**
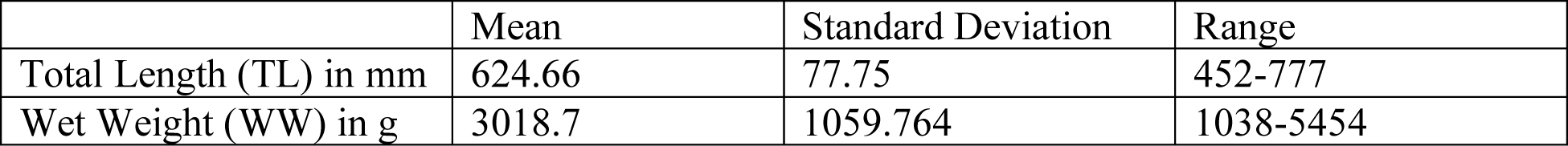
Total length and wet weight for 50 acoustically tagged silver carp sampled in the James River in Summer 2021. TL is in mm, and WW is in grams.

|  | Mean | Standard Deviation | Range |
| --- | --- | --- | --- |
| Total Length (TL) in mm | 624.66 | 77.75 | 452-777 |
| Wet Weight (WW) in g | 3018.7 | 1059.764 | 1038-5454 |

**Table 2:** Model comparisons of each movement model using WAIC (Hooten & Hobbs, 2015). The best fit model is in the top row and the least best fitting model in the last row.

| Model parameters | elpd_diff | se_diff |
| --- | --- | --- |
| Discharge | 0.0 | 0.0 |
| Discharge, Δdischarge24h | -0.1 | 0.1 |
| Discharge, Δdischarge48h | -0.2 | 0.1 |
| Discharge, weekly mean DO | -0.4 | 0.5 |
| Discharge, weekly mean temp | -0.6 | 0.3 |
| Discharge, Δdischarge24h, weekly mean temp, weekly mean DO | -0.6 | 0.5 |
| Discharge, weekly mean DO, weekly mean temp | -0.7 | 0.5 |
| Discharge, Δdischarge48h, weekly mean DO | -0.7 | 0.5 |
| Discharge, Δdischarge24h, weekly mean temp | -0.7 | 0.4 |
| Discharge, Δdischarge48h, weekly mean temp, weekly mean DO | -0.7 | 0.5 |
| Discharge, Δdischarge24h, mean weekly DO | -0.7 | 0.5 |
| Discharge, Δdischarge48h, weekly mean temp | -0.8 | 0.4 |
| Weekly mean temp, weekly mean DO | -1.7 | 1.3 |
| Δdischarge48h | -2.1 | 1.4 |
| Weekly mean temp | -2.4 | 1.6 |
| Δdischarge24h | -2.5 | 1.5 |
| Weekly mean DO | -3.6 | 1.8 |

The mean distance moved by all tagged individuals (excluding fish of “undetermined” fate) during the entire study period (June 2021 to October 2022) was 363 rkm (range: 0 to 1385.1 rkm, Figure 2). High discharge within the James River from May through June of 2022 (Figure 3a) caused most tagged individuals to undergo their highest recorded movement proportions of the entire study period (Figure 3b). The most upstream silver carp detection was near Huron, SD (rkm 354), while the most downstream detection was just outside of Blair, NE, in the Missouri River (rkm -236). Only one of the tagged individuals was recorded swimming out of the James River (and returned to the James River shortly thereafter), and all other individuals remained within the James River throughout the study period (Figure 4).

**Figure 2.**
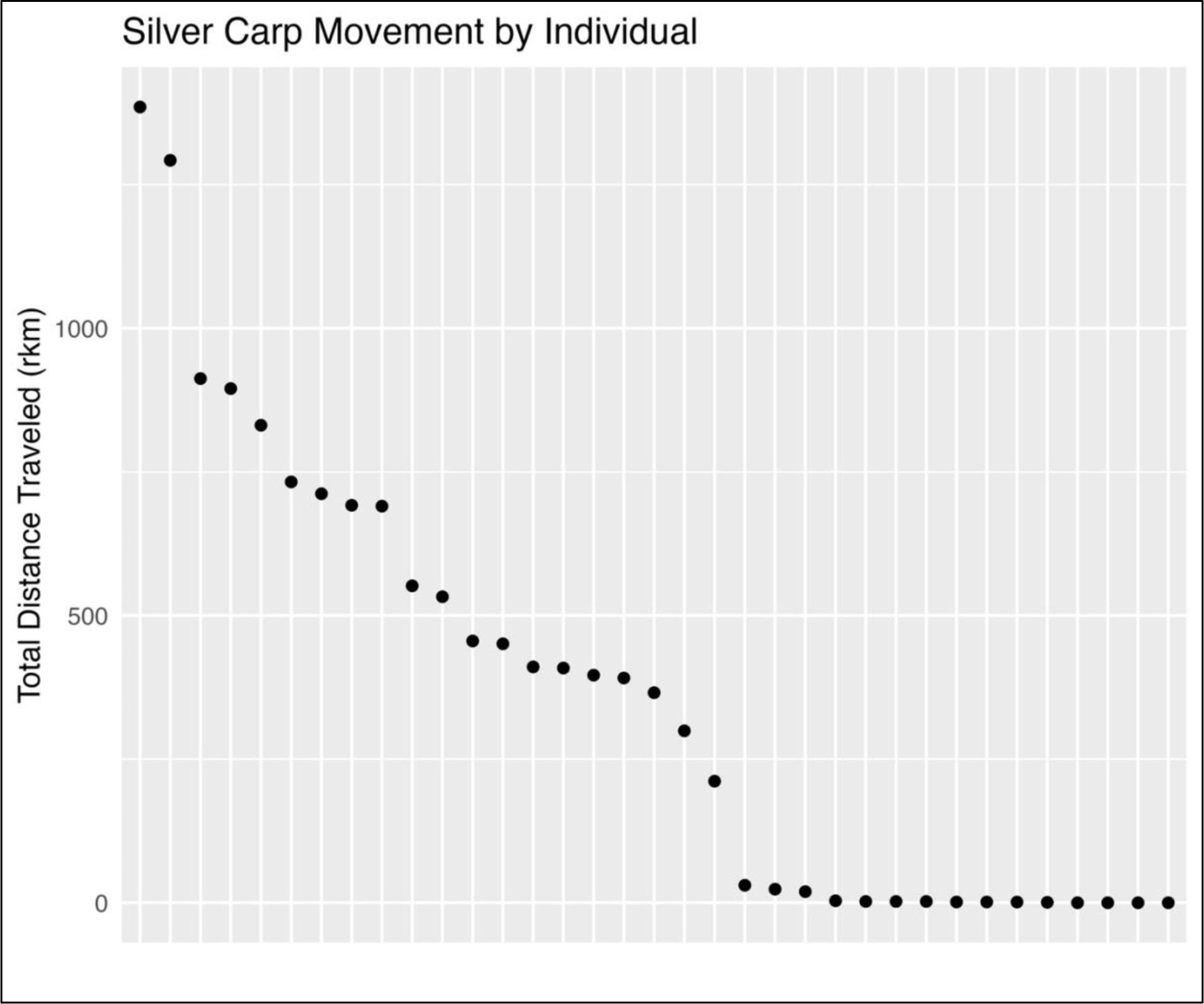
The total distance traveled (in river kilometers) by telemetered silver carp in the James River, SD from June 24, 2021 to October, 24 2022. Each black dot represents one telemetered individual. The mean total distance was 363 rkm (range: 0 to 1385.1 rkm). The 15 fish with “undetermined” fate were not included.

**Figure 3.**
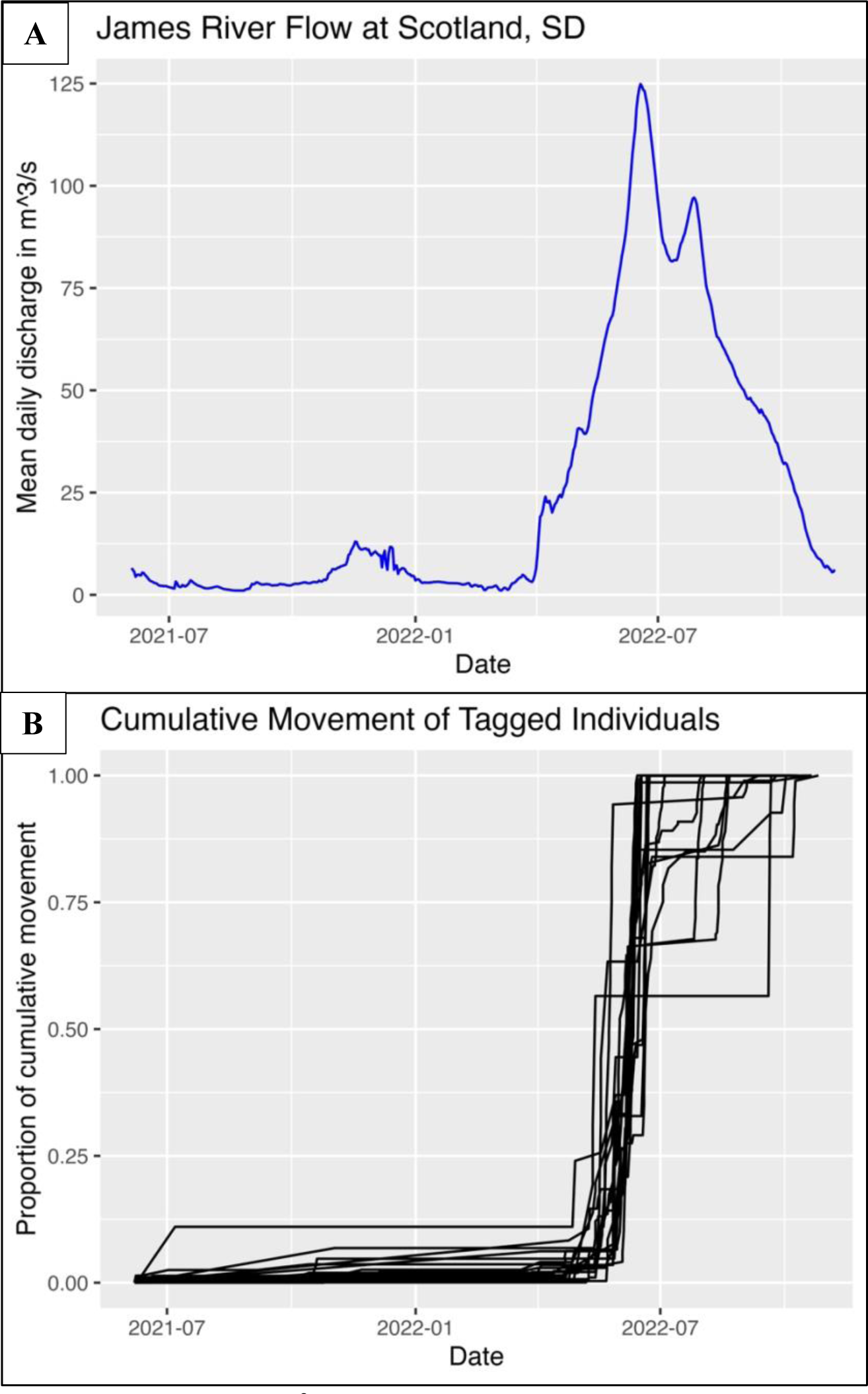
a) Average daily discharge (m^3^/sec) in the James River; data pulled from the USGS gage station # 06478500 (James River near Scotland, SD) using the *dataRetrieval* package (De Cicco and Hirsch 2014) and visualized using *ggplot2* (Wickham et al. 2016); 3b) Cumulative movement over time for telemetered silver carp on a scale from 0 to 1, with 0 representing no movement and 1 representing the cumulative total of each individual’s movement.

**Figure 4.**
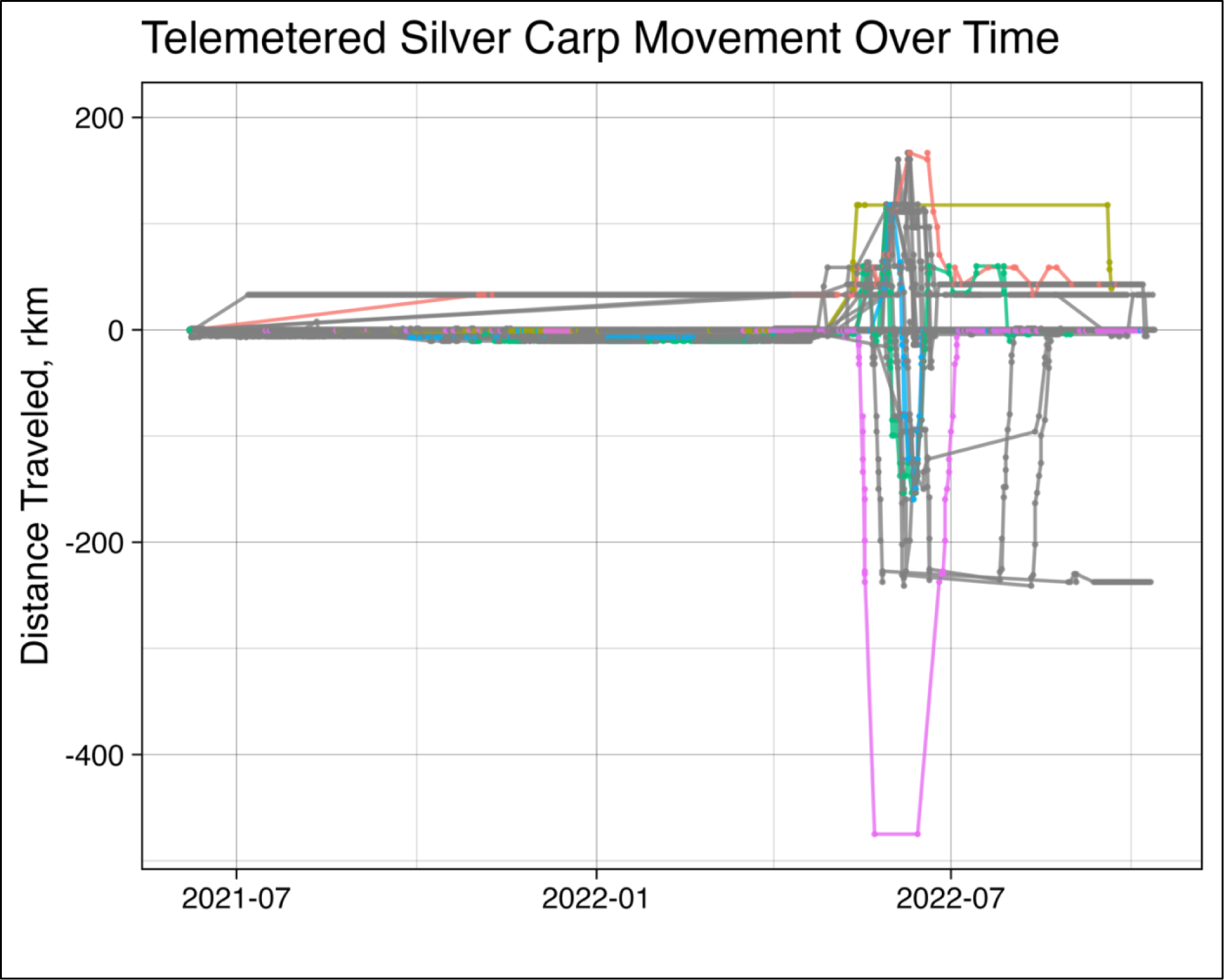
Movement data relative to the tagging location for silver carp in the James River, South Dakota. Zero on the y-axis represents the tagging location for each fish. Negative river kilometers on the y-axis represent downstream movement from the initial tagging location, and positive river kilometers represent upstream movement relative to the initial tagging location. A few individuals are colorized to show different variations in movement patterns.

During this study, daily mean flow ranged from 1.4 to 124.9 m^3^/s, change in discharge over 24 hours ranged from -5.1 to 5.1 m^3^/s, change in discharge over 48 hours ranged from -6.5 to 9.3 m^3^/s, temperature ranged from 0 to 31°C, and dissolved oxygen ranged from 20 to 201% O2.

Of all predictor variables modeled in this study (daily mean discharge, change in discharge over 24 hours, change in discharge over 48 hours, mean weekly dissolved oxygen, mean weekly temperature), the best fitting model to predict silver carp movement included only daily mean discharge (flow) as a predictor. According to the best fitting model, when flow was low (1.5 m^3^/s), there was a 59% probability that silver carp would move (95% CrI: 34% to 81%). The probability of movement increased as flow increased, with a 63% probability (95% CrI: 40 to 83%) at 8.5 m^3^/s, 74% probability (95% CrI: 57 to 87%) at 30 m^3^/s, and 94% probability (95% CrI: 80 to 99%) at 100 m^3^/s (Figure 5).

**Figure 5.**
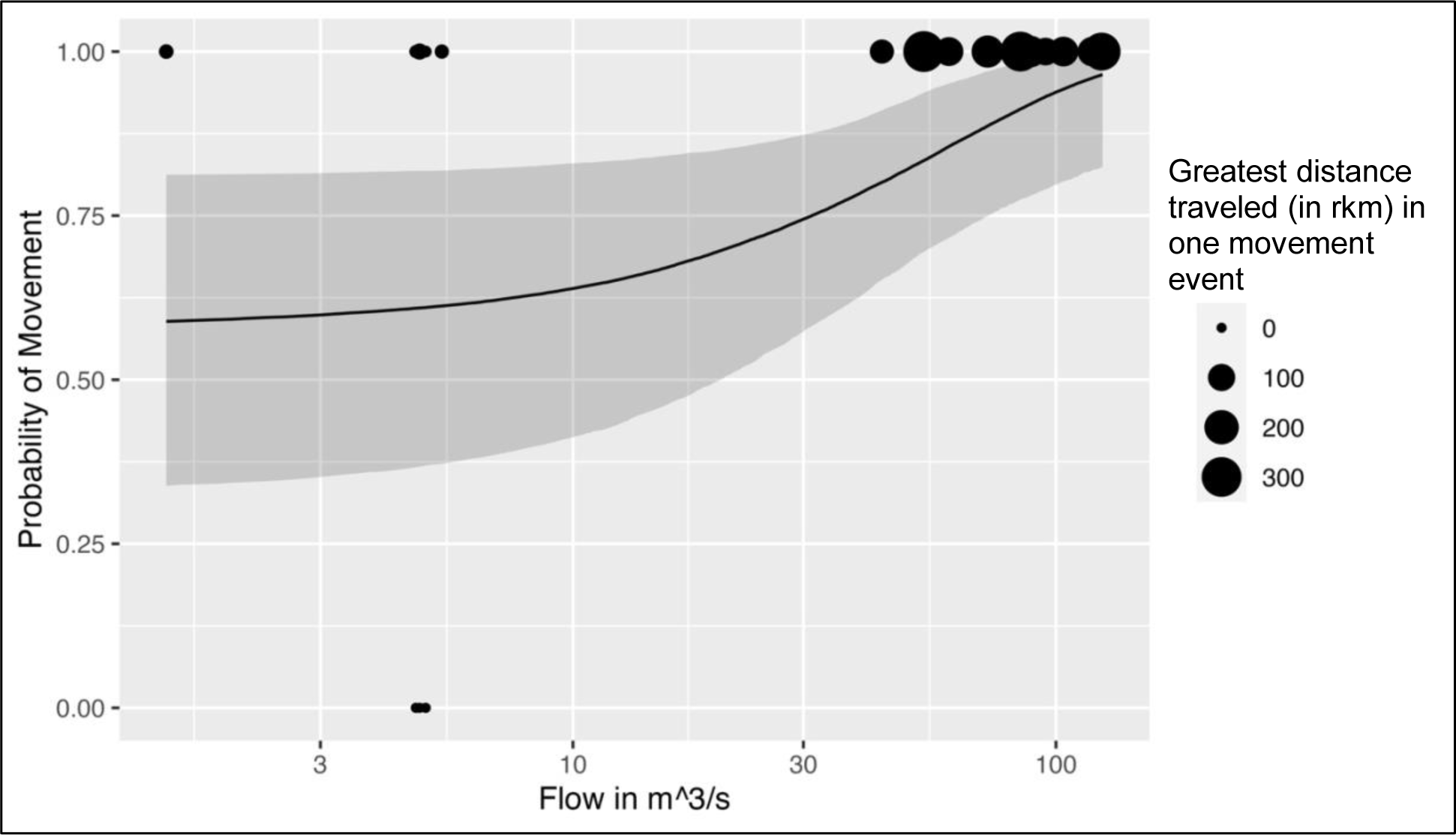
Posterior prediction intervals for the probability of movement as flow increases. Dots represent each individual and are scaled to represent the greatest distance the individual traveled in one movement event. Most individuals who moved during high flows also moved distances of over 200rkm in one movement event.

### Model performance

All models performed well, and observed data was within the predicted range of the posterior predictions. Trace plots indicated proper mixing of the Markov chains, all R-hat values were < 1.02, and all parameter estimates had large effective sample sizes. The Bayesian R^2^ value for the best fitting model was 0.2019 (95% credible interval (CrI): 0.0087 to 0.4171). Posterior predictive checks (Berkhof et al. 2000), generally replicated the shape of the data for all models. The sensitivity analysis showed model parameter estimates were not substantially changed from the original best fitting model parameter estimates.

## Discussion

The most important result of this study is that average daily discharge (flow) was the best predictor for silver carp movement in the James River, South Dakota. As mean daily flow increased, the probability of silver carp movement also increased. Movement data also suggests that the tagged individuals were resident within the James River. Even when the single tagged fish left the James River, it returned to almost the exact location that it had previously been occupying in the James River after a short period of time.

Understanding cues to movement and observing broad-scale movement patterns is an important first step in limiting the spread of invasive species. In our study, we recorded some sedentary individuals (total movement <50 rkm (n=17)) and some active individuals (total movement >50 rkm (n=18). This is a common occurrence within freshwater fish populations (Leggett 1977; Rodríguez 2002) and moreover, is common in other populations of telemetered silver carp in places such as the Wabash River in Illinois (Prechtel et al. 2018; Coulter et al. 2022). The propensity for movement for each individual fish can depend upon factors such as availability of food resources (Olsson et al. 2006), seasonality (Brodersen et al. 2008; Brönmark et al. 2014), personality traits such as “boldness” (Chapman et al. 2011), and phenotypic traits (Skov et al. 2011). For silver carp specifically, factors influencing movement propensity include increases in temperature and discharge which trigger spawning movements, movements upstream or downstream to colonize new habitats, and behavioral plasticity leading to greater movement ability and increased invasion success. (Coulter et al. 2016; Coulter et al. 2018; Coulter et al. 2022; Lebsock et al. 2020). Because the tagged individuals in our study almost always returned to their tagging locations, home range may also play a major role in the movement ecology of these carp in the James River.

Average daily discharge (“flow”) appeared to have the greatest influence on predicting silver carp movement in the James River. This result parallels that of other silver carp movement studies. (Coulter et al. 2022; Lebsock et al. 2020; Prechtel et al. 2018). Tagged silver carp within the James River remained mostly stationary until the mean daily discharge reached ∼42 m^3^/s.

This provides a target range of discharge values (i.e. 0-42 m^3^/s) during which population control efforts could be implemented to maximize removal success. Incidentally, we observed that silver carp tended to school in deeper habitats, such as within river bends and in deep backwater areas. However, the use of side-scan sonar or other similar imaging methods could more clearly elucidate habitat preference and further pinpoint preferred habitats to increase the efficiency of removal efforts. In the James River, if targeted removal were to be performed in areas with known silver carp congregations during times of low flow, it may be possible to maximize removal success while minimizing the labor and costs associated with removal efforts.

Continuing to collect telemetry data to encompass a wider range of environmental conditions, while also including active tracking data and side-scan sonar, would provide a picture of finer-scale carp movement in areas where the receiver network is lacking, and would allow for a deeper understanding of fine scale movement patterns and habitat use preferences.

There are a few caveats to this study. The receivers within our telemetry array did not overlap in detection range and therefore the recorded distances could only provide a picture of large-scale movement patterns of silver carp in the James River. We had multiple fish that seemed to “disappear” from our network and then reappear on the same receiver at a later date (n=4), but overall net movement was still recorded as zero for that individual. We also had 15 fish that were never detected on a receiver after their initial tagging date. Increasing the number of receivers and decreasing the distance between receivers could provide data for finer-scale movement patterns. We were also only able to collect 1.5 years of telemetry data, which limits our interpretation of movement data results to the time scale and environmental conditions we observed during the study period. The years had vastly different discharge (2021 was a “low-water” year and 2022 was a “high-water” year for the James River (Carlson 2022)) and the individuals with the most movement in 2022 were often sedentary in 2021. Extending the study for a longer period would encompass more environmental conditions and improve the overall interpretation of telemetry results. While we were unable to assign sex during tagging, sex could be a valuable variable to include in future studies, as sex and sex-dependent traits determine movement patterns in many other species of freshwater Cyprinid fishes (Lucas and Batley 1996).

## Conclusion

The James River in eastern South Dakota is on the northwestern-most leading edge of the silver carp invasion. Their movement patterns and habitat use were not previously studied in this system. We found that individuals exhibited a wide range of total movement during the study period (mean: 256.7 river kilometers; range: 0 to 1385.1 river kilometers). As daily mean discharge reached ∼41 m^3^/s, there was an 80% probability that silver carp would move (95% CrI: 64 to 90%). Understanding movement dynamics on this side of the invasion front is key to preventing further northwestward expansion of the species into habitats such as the major reservoirs on the mainstem Missouri River. This study provides targets for informed management of the species and suggests that removal efforts should take place when mean daily discharge in the James River is below 41 m^3^/s to maximize removal success.

## Acknowledgements

We thank B Schall, T Carlson, T DeVine, J Larsen, J Stahl, N Loecker, K Schwartz, and the SDGFP 2021 Summer interns for their assistance with fish collection and surgeries, deploying receivers, and retrieving data. Funding for this study was provided South Dakota Game Fish and Parks (Grant number F-16-R-1) and the United States Fish and Wildlife Service (Grant number F20AP11067-00). This work was conducted under an approved protocol from the University of South Dakota’s Institutional Animal Care and Use Committee (protocol # 10-08- 20-23D).

