## Supplemental Figures for "Using passive telemetry and environmental variables to predict Silver Carp (*Hypophthalmichthys molitrix)* movement cues on the northwestern edge of their invasion front"

**Supplemental Figures**

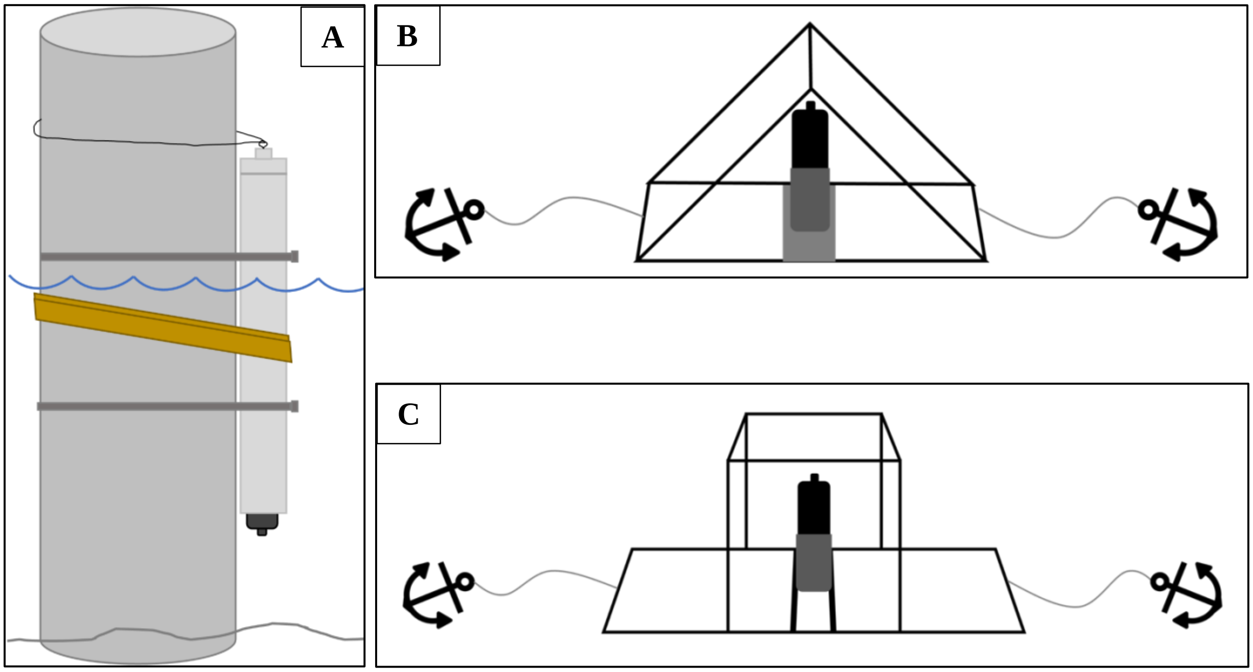


Figure S1a) A sketch of the bridge-mounted receiver setup. The PVC pipe is secured using two metal bands (grey), a double-wrapped tow strap (dark yellow). The receiver is suspended using two lengths of 6mm wire within the PVC pipe and is exposed out of the bottom of the PVC pipe by at least 5 cm. Picture adapted from the drawings in Carlson (2022) and Carlson et al., (2023) with permission; S1b) Drawing of the triangle-shaped frame receiver mount; S1c) Drawing of the square-shaped frame receiver mount.

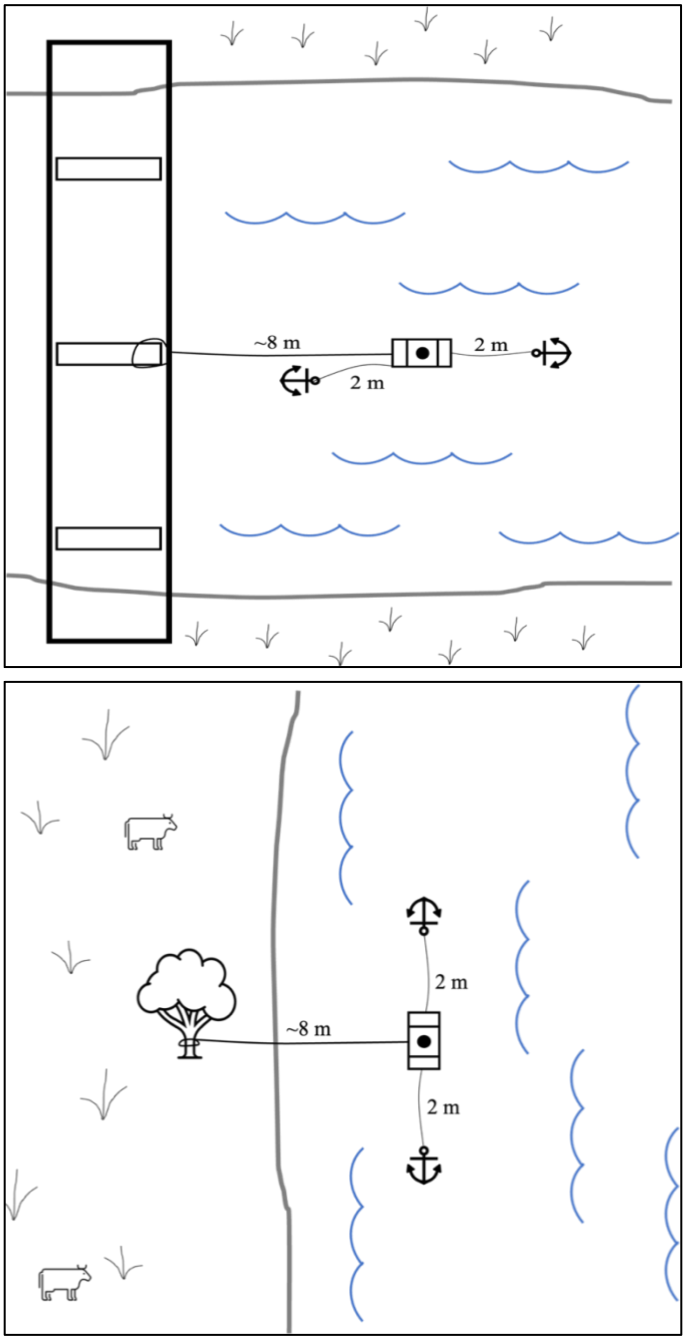


Figure S2a) A top-down schematic of the frame mounted receiver attached to a bridge with metal crossbar supports; S2b) A top-down schematic of the frame mounted receiver attached to a tree on the bank, adapted from the drawings in Carlson (2022) and Carlson et al., (2023) with permission.
